# Tackling Anticancer Drug Resistance and Endosomal Escape in Aggressive Brain Tumors Using Bioelectronics

**DOI:** 10.1101/2024.06.03.597127

**Authors:** Akhil Jain, Philippa Wade, Snow Stolnik, Alistair N. Hume, Ian D. Kerr, Beth Coyle, Frankie Rawson

## Abstract

Chemotherapy resistance and endosomal entrapment, controlled by intracellular trafficking processes, are major factor in treatment failure. Here, we test the hypothesis that external electrical stimulus can be used to modulate intracellular trafficking of chemotherapeutic drugs in most common malignant brain tumors in childhood (medulloblastoma) and gold nanoparticles (GNPs) in adulthood (glioblastoma). We demonstrate that application of alternating current (AC) with frequencies ranging from KHz-MHz and low strength (1 V/cm) lead to killing of cisplatin and vincristine resistant (mediated by extracellular vesicles) medulloblastoma cell lines. On the other hand, in primary glioblastoma cells high frequency AC (MHz) regulated the endosomal escape of GNPs. No significant effect on the viability of the control medulloblastoma cells (resistant cells cultured in drug free media and non-resistant cells) and glioblastoma cells after AC treatment confirmed targeting of intracellular trafficking process. This work supports future application of AC in drug delivery and brain cancer therapy.

## 1. Introduction

Cells have honed their ability to maintain homeostasis through delicately balancing their internal environment by exerting precise control over what enters and exits the cell. This fine-tuned regulation is primarily orchestrated by cell and organelle membranes, the gateway to the cell and organelles, respectively.^1^ The implications of this cellular control mechanism extend far beyond basic biological processes. They hold profound significance in the realm of disease treatment, particularly in the context of combating cancer. Cancer cells are notorious for their ability to evade therapeutic interventions and have evolved strategies to resist the effects of chemotherapy drugs through membrane bound organelles and particles such as endosomes and extracellular vesicles (EVs), respectively effectively thwarting their intended actions.^2^ This adaptation highlights the critical role played by cellular homeostasis in the development of resistance and underscores the need for novel approaches to tune these transport mechanisms.

EVs play a crucial role in the function of glioblastoma (GBM) and more broadly gliomas, which are difficult-to-treat cancers.^3,4^ Within this context, EVs have gained prominence due to their critical role in various cellular processes, including intercellular communication, cell signaling, and immune regulation.^5^ GBM cells secrete these small membranous vesicles, containing an array of cargo molecules such as proteins, RNA, and lipids.^6^ EVs play a significant role in GBM biology and pathogenesis, where they contribute to tumor growth, invasion, and metastasis by delivering oncogenic proteins and signaling molecules to other cells in the tumor microenvironment, suppressing the anti-tumor immune response, promoting angiogenesis and inducing resistance to chemotherapy and radiation therapy^5^.

EVs have also emerged as crucial transport system in medulloblastoma. Medulloblastoma-derived EVs promote tumor growth, invasion, and metastasis, often by mechanisms akin to those observed in GBM-derived EVs.^7^ The therapeutic potential of EVs in GBM and medulloblastoma is under exploration. EVs could serve as vehicles for delivering therapeutic drugs or genes to tumor cells, modulating the tumor microenvironment to suppress tumor growth and progression, or developing vaccines targeting tumor-specific antigens.^8^ Furthermore, the role of EVs as biomarkers for monitoring disease progression and therapy response in patients with these aggressive brain tumors holds great promise.^9^ Importantly for this study, the reviews of the involvement of EVs in gliomas reveal evidence suggesting that glioma cells utilise the vesicles to expel therapeutics, thereby enhancing resistance.^4^ The challenges encountered in advanced drug delivery further underscore the importance of understanding cellular homeostasis.^10^ A long standing challenge in advanced drug delivery is that drugs and nanoparticles can be trafficked and siloed in endosomes and subsequently degraded in lysosomes. To date, only a small fraction of these systems has advanced to clinical use, mainly due to issues such as entrapment in endosomes and degradation in lysosomes.^11^ For instance: delivery of siRNA and other chemotherapeutics by lipid nanoparticles or EV based therapeutic delivery is limited by endosomal entrapment.^12, 13^ This unfortunate fate can render potentially life-saving medications ineffective.^12^

Consequently, there is an urgent need to develop widely applicable technologies that can modulate and precisely control the trafficking of drug, molecules, and nanoparticles within cells. By unraveling the intricate mechanisms and developing disruptive technological approaches to govern intracellular transport, we can begin to overcome the challenges posed to drug delivery and enhance the efficacy of treatments. Such advancements hold promise for revolutionising the field of medicine and could one day improve patient outcomes.

Bioelectronic medicines are an emerging therapeutic approach in which electrical input can be used for the treatment of disease. There has been a recent shift to develop wireless electrical system for triggering release of drugs from their carriers, as reviewed by Mirvakili and Langer ^15, 16^. Additionally, electroporation has been heavily studied for enhancing cell delivery of an anti-cancer drug with the most recent study investigating this in single cells.^17^ We have previously demonstrated that ultrasound can be used to intracellularly release drug from liposomes.^18^ Our group has recently shown that *AC* could bypass the plasma membrane to exert effects across the cell membrane.^16, 19^ Moreover, it has been established that electrical input on lipid bilayers causes subtle structural perturbations in their structure.^20^ Therefore, we hypothesized that by exploiting electrical input intracellular fate of nanoparticle-based delivery systems for anti-cancer therapy could be modulated. This could influence two distinct processes (i) in cancer treatment: transport of anticancer drug out of cell via EVs; moderating EVs transport processes would lead to increase cytoplasmic exposure of the drug and cancer cells killing, and (ii) Release of GNPs from endosomal/lysosomal compartment into cytoplasm. This could lead to generation of new nanomedicine for improved therapeutic outcomes. To the best of our knowledge, there has been no demonstration of using AC to prevent transport of chemotherapies outside of cells via EVs. Furthermore, the added potential of using AC would increase the drug delivery systems presence within the cytoplasm by facilitating endosomal escape.

In this work, we conducted a study aiming to merge electronics for delivery of AC with chemotherapy to chemo-resistant (Cisplatin - cis and Vincristine – vin) medulloblastoma and GBM cells. Our objective was to investigate whether this delivery of High frequency-AC (HF- AC) to cells could help overcome the problems of (i) EV-mediated drug transport out of cancer cells and (ii) modulate the intracellular escape of nanoparticles from endosomes by influencing transport process. The results of our study successfully demonstrate this concept, indicating that by combining electronics with anticancer therapy, we can potentially introduce a new bioelectronic technology that holds promise for the treatment of resistant cancers. Further development of this approach could enhance the efficacy of chemotherapies and more broadly the field of cancer therapeutics.

## 2. Methodology

### 2.1. Cell lines and standard culture conditions

Nomenclature and identifiers of the cell lines used in this work is as follows: DT = drug tolerant, ^DT^ = drug treated, and ^NDT^= non-drug treated. For e.g. DT-XXX-Cis/Vin^DT^ means drug tolerant (DT) XXX cell line, tolerant to Cis or Vin and are drug treated (^DT^). Similarly, DT-XXX-Cis/Vin^NDT^ means drug tolerant (DT) XXX cell line, tolerant to Cis or Vin and are non-drug treated (^NDT^). Vehicle cells are a matched DMSO or DMF vehicle cell line was generated alongside to account for morphological and genetic changes resulting from vehicle (DMF or DMSO) exposure and long-term culture. For e.g. DT- XXX-DMSO/ DMF = Drug tolerant (DT) XXX vehicle cell line grown in cell culture medium containing equivalent volume of DMSO or DMF that has been used in drug treated tolerant lines.

Sonic-hedgehog (SHH) medulloblastoma DAOY cell line was purchased from ATCC (ATCC® HTB-186™) and grown in DMEM with 10% fetal bovine serum (FBS, HyClone (Logan, Utah, USA). Three vehicle and cis-tolerant MB Group3 cell lines (DT-D283-DMF, DT-D283-Cis, DT-HD-MB03-DMF, DT-HD-MB03-Cis, DT-D458-DMF and DT-D458-Cis) were utilized, as previously published.^21^ The DT-D283 and DT-D458 lines were derived in-house whereas the DT-HD-MB03-Cis was obtained from Gianpiero Di Leva (Keele University, UK). DT-D283 and DT-D458 cells were cultured in DMEM with 10% fetal bovine serum and the DT-D283-Cis^DT^ and DT-D458-Cis^DT^ supplemented with 1.6 and 0.6 µM cis (Selleckchem, Houston, TX, USA, S1166) respectively. DT-HD-MB03 were grown in RPMI 1640 with 10% FBS and the DT-HD-MB03-Cis^DT^ cells were supplemented with 0.5 µm cis. The equivalent volume of vehicle (DMF) was added to the matched vehicle line. All cell lines were mycoplasma tested monthly and grown in antibiotic-free culture conditions at 5% CO2 and 37 °C. EV-depleted FBS was generated by ultracentrifugation at 100,000 × *g* at 4°C for 18 hours. Filter sterilization using a 0.22 µm filter (Millipore) was carried out prior to its addition to culture medium, resulting in EV-depleted medium. GBM cells - Glioma INvasive Marginal 31 (GIN) cells from the infiltrative tumour margin and Glioma Core Enhanced 31 (GCE) from the core of the tumor were isolated previously. ^22,16^ Both GIN and GCE cells were cultured in DMEM (Gibco) supplemented with 10% FBS, 1% Penicillin/Streptomycin and 1% L-Glutamine. Cells were maintained at 37°C in an incubator, containing 5% CO2. Cells were tested for mycoplasma every month, where they were grown in an antibiotic-free medium for one week before mycoplasma testing. All cells used were mycoplasma-free.

### 2.2 Generation of vincristine- and cisplatin-resistant cell lines

A continuous model of selection was used to generate drug-tolerant MB cell lines resistant to vin and cis. SHH DAOY cells were cultured continuously in the presence of vin (Selleckchem, S1241) and the concentration dose was escalated upon cell proliferation. Cells were passaged in T-25 flasks and initially treated with 1/10th of their vin EC50 and the dose increased upon cell proliferation. A matched DMSO vehicle cell line was generated alongside to account for morphological and genetic changes resulting from vehicle exposure and long-term culture. Cells were considered resistant when the EC50 against vincristine for the treated cells had exceeded the treatment dose and was significantly increased in comparison to the EC50 of the vehicle line. DT-DAOY cells were grown in DMEM with 10% FBS, and the DT-DAOY-Vin^DT^ line supplemented with 2.8 nM vin. Akin to the DT-DAOY cells, cis-resistant cell lines DT- D458, DT-HD-MB03 and DT-D283 were also generated in a similar manner.^21^ In brief, cells were treated with increasing doses of cis until their EC50 exceeded the treatment dose or was significantly increased in comparison to the vehicle cell line (DMF).

### 2.3 Drug cytotoxicity assay

Cells were seeded into clear bottomed, black-walled 96-well plates (Greiner; 655096) at a density of either 1 ×10^3^ (DT-DAOY), 5 × 10^3^ (DT-HD-MB03) or 1 × 10^4^ (DT-D283 and DT-D458) cells per well and left overnight. Cells were then challenged with varying concentrations of vin or cis prior to being incubated for 72 hours at 37°C and 5% CO2. After 72 hours, metabolic activity was assessed using PrestoBlue (Thermo Fisher; A13262) and fluorescence was measured using the FLUOstar Omega microplate reader at 560/590 nm. Cell viability was calculated as a percentage relative to the vehicle control and EC50 were calculated in GraphPad PRISM 9 using nonlinear regression with three parameters. Significant differences between EC50 were calculated using one-way ANOVA with Sidak’s multiple comparisons. The data represents the SEM of three independent experiments.

### 2.4 Isolation of extracellular vesicles

To isolate extracellular vesicles from cell cultures, cells were grown in 1 × T-225 flask up to 30% confluence, washed twice with Hanks’ Balanced Salt Solution (HBSS, Gibco (Loughborough, UK)) and incubated in EV-depleted medium for 48h. Cell culture medium was collected and centrifuged at 300 × *g* for 5 minutes to pellet cells. The supernatant was then centrifuged again at 1,500 × *g* for 10 minutes, followed by a final centrifugation at 10,000 x *g* and 4°C for 10 minutes to remove any debris and large particles. The supernatant was filtered through a 0.22 µm filter prior to ultrafiltration using a 100K MWCO protein concentrator (Thermo Scientific™ Pierce™; 88533) where the supernatant was centrifuged at 3,000 × *g* until a ∼ 1 mL concentrate was left. The 1 mL concentrate was loaded directly onto size-exclusion chromatography columns obtained from Izon (qEV1 / 70 nm; IC1-70) and EV fractions collected according to manufacturer’s instructions. Concentration of EVs was determined using ZetaView® Nanoparticle Tracking Analysis relative to 1 × 10^6^ cells.

### 2.4. AC Stimulation

Electrical Stimulation (ES) with AC was carried out by inserting two steel electrodes (0.5 mm × 25 mm) at the opposite end (fixed at 10 mm from each other) of each well in a 24-well plate and dipped in cell culture medium. These electrodes were connected to an Arbitrary Function Generator (AFG-21225, RS PRO, UK) which delivered the AC sine-wave signals, frequency, and amplitude. The cells were stimulated with AC with a desired frequency and peak voltage amplitude of 1 V/cm for 30 minutes. The strength of AC between the electrodes was measured using a digital oscilloscope (TDS 210, Tektronix).

### 2.3. Metabolic activity assay

The medulloblastoma vehicle cell lines DT-D283-DMF, DT- D458-DMF, DT-HD-MB03-DMF and DT-DAOY-DMSO and the cell lines resistant to cis - drug treated (DT-D283-Cis^DT^, DT-D458-Cis^DT^, and DT-HD-MB03-Cis^DT^) and non-drug treated (DT-D283-Cis^NDT^, DT-D458-Cis^NDT^, and DT-HD-MB03-Cis^NDT^) and vincristine - drug treated (DT-DAOY-Vin ^DT^) and non-drug treated (DT-DAOY-Vin^NDT^) were seeded at density of 1.0 × 10^5^ per well in a 24-well plate. The cells were stimulated with AC using the protocol mentioned above in section 2.4. Immediately after stimulation with AC the cells were incubated at 37°C and 5% CO2 for 24 h before carrying out metabolic activity assay. Next, the medium containing cells was replaced with fresh medium containing 1% PrestoBlue^TM^ HS cell viability reagent (ThermoFisher Scientific, UK) and incubated for 1 hour before reading the fluorescence at 590 nm/610 nm (excitation/ emission) in a microplate reader (Tecan Infinite M Plex and Spark 10M). Cells grown in culture medium without AC treatment were used as the negative control. Results are presented relative to negative control. The data is represented as an average of triplicate experiment with 3 independent repeats ± S.E.M.

### 2.4. Live/Dead assay

After stimulation with AC the cells were incubated for 24 h in an incubator at 37°C and 5% CO2. Next, cis resistant and vehicle cells were centrifuged at 300 g for 5 minutes (due to their semi-adherent nature), and the pellet was dispersed in fresh medium containing 1 mM Calcein AM and 1 mg/mL Propidium iodide (ThermoFisher, UK), and incubated for 30 min at 37°C and 5% CO2 in a 24 well plate. The cells were then centrifuged (300 g for 5 minutes), and the cell pellet was washed with PBS. Finally, the cells were placed in a 24-well plate (m-plate 24 well black, ibiTreat, Thistle Scientific, UK) in phenol red free medium and imaged using a Leica TCS SPE Confocal Microscope. The proportions of live and dead cells were quantified using ImageJ software.

### 2.5. Biocompatibility of gold nanoparticles

The GIN and GCE cells were seeded in a 96-well plate at a density of 5 × 10^3^ cells/well and allowed to adhere for 24 h. The medium was replaced with fresh medium containing Texas red conjugated 100 nm spherical gold nanoparticle (GNP, Nanopartz, Inc, USA) at different concentrations (25, 50, and 100 μg/mL) and the cells were incubated for 8 h. Cells with nanoparticle suspension were then stimulated with AC of various frequencies at 1 V/cm for 30 min. Following stimulation cells were then incubated for 24 h at 37°C and 5% CO2. Next, the medium was replaced with complete medium containing 10% PrestoBlue^TM^ HS cell viability reagent (ThermoFisher Scientific, UK) and incubated for 1 h before reading the fluorescence at 590 nm / 610 nm (excitation/ emission) in a microplate reader (Tecan Infinite M Plex and Spark 10M). The data is represented as an average of triplicate experiment with 3 independent repeats.

### 2.6 Endo/lysosomal escape

GIN 31 and GCE 31 cells were seeded at a density of 4 × 10^4^ cells per well in a 24-well plate and incubated at 37°C and 5% CO2. After 24 h, the culture medium was replaced with fresh medium containing CellLight™ Late Endosomes-GFP, BacMam 2.0 (ThermoFisher Scientific, UK) and incubated overnight at 37°C and 5% CO2. Later, the medium was replaced with fresh medium containing 25 μg/mL of GNP (in PBS) and incubated for 8 h. Immediately after the ES, the cells were washed with PBS and imaged using a Leica TCS SPE Confocal Microscope.

## 3. Results and Discussions

### 3.1. Drug resistance in medulloblastomas

Cis and Vin are two of the standard of care chemotherapies for medulloblastoma. Previously we have described three cis resistant medulloblastoma cell lines;^21^ in that work, compared to matched vehicle controls, DT-D458-Cis^DT^ showed 8-fold resistance, DT-HB-MB03-Cis^DT^ showed 2-fold resistance and DT-D283- Cis^DT^ showed 5-fold resistance (for cell line identifier please refer to methodology, section 2.1.). In the current work (Fig. 1) we established vin tolerant DAOY cell line (DT-DAOY- Vin^DT^) by continuous treatment of cell culture with escalating concentrations of vincristine. DT-DAOY-Vin^DT^ has a 4-fold increase in the EC50 compared to the matched vehicle cell line (DT-DAOY-DMSO) (p<0.001, Fig. 1).

**Figure 1.**
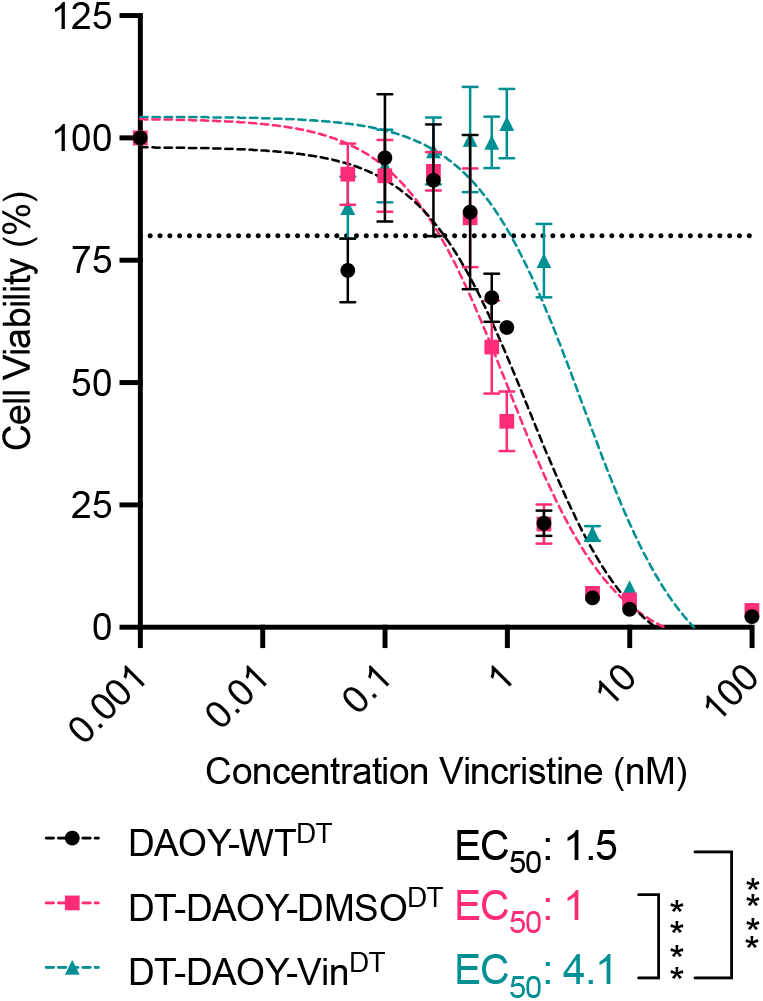
Continuous long-term vincristine treatment promotes increased cell resistance to vincristine in medulloblastoma SHH DAOY cells. The cells were initially treated with vin at 1/10^th^ of their EC50 and upon cell proliferation, the cells were subsequently challenged with an increasing dose of vin. Cell viability was assessed via drug-response assays and the EC50 calculated using nonlinear regression analysis with three parameters. For detailed description of the cell lines and their identifiers used in this work please refer to methodology section 2.1. DAOY-WT^DT^ = Wild-type DAOY cell line treated with Vin, DT-DAOY-Vin^DT^ = Vin tolerant cell line treated with Vin and DT-DAOY-DMSO^DT^ = Vehicle cell line treated with Vin. Significance was assessed using one-way ANOVA with Šídák’s multiple comparisons test. ****p ≤ 0.0001.

Intriguingly, comparison of the number of EVs released revealed a significant increase by our cis tolerant lines (Fig. 2 a-c) relative to their vehicle counterpart (Fig. 2). On the other hand, in vin tolerant lines no significant difference was observed compared vehicles cells (Fig. 2d). Nevertheless, an increase in average number EVs in drug treated vin tolerant lines was observed, which can be further supported with literature, where it has also indicated that EVs can act as transporters for efflux of drugs.^23^ Thus it can be concluded that number of EVs released is enhanced in drug tolerant lines which together with EC50 data suggests the EV mediated drug tolerance in medulloblastoma cell lines.

**Figure 2.**
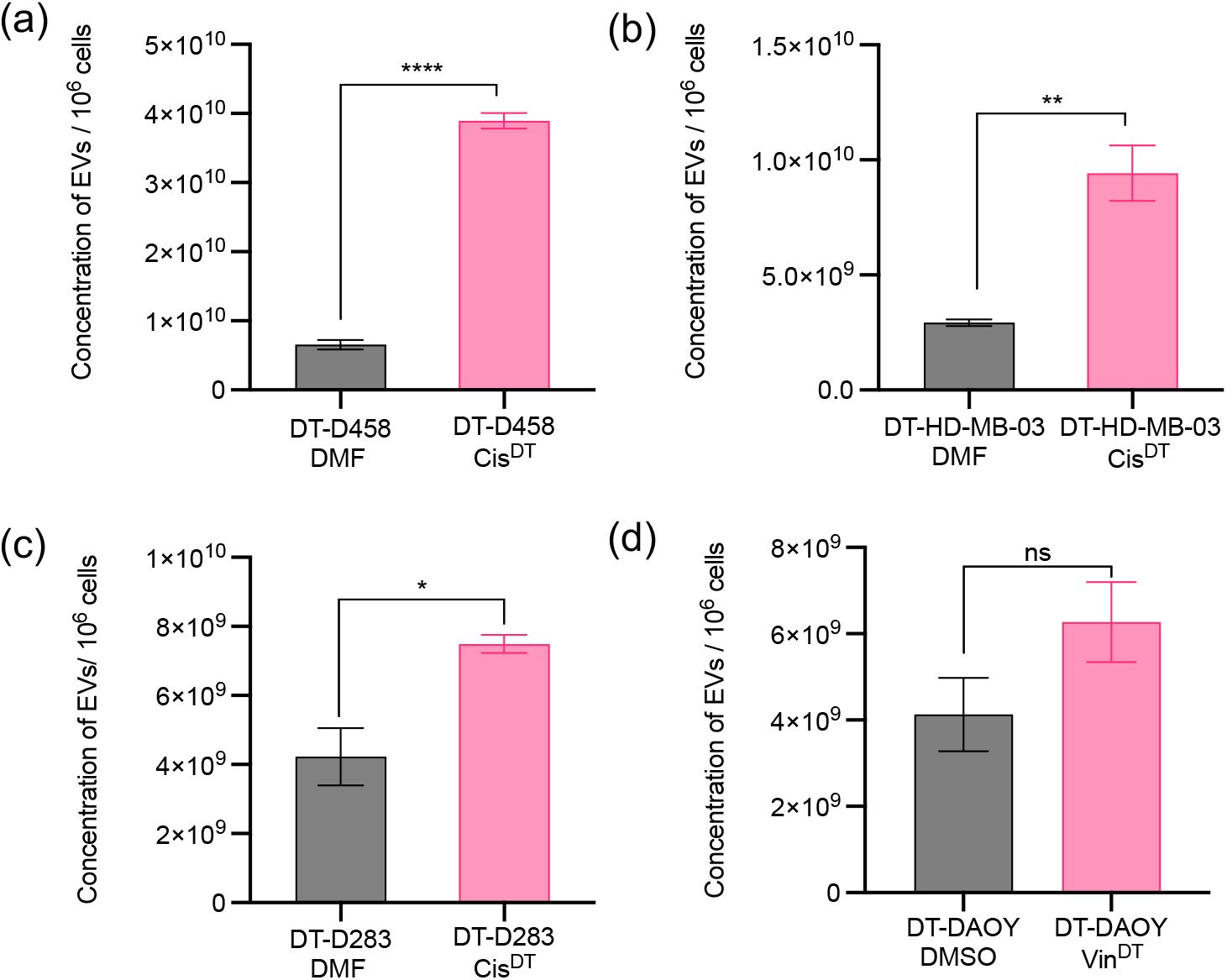
Quantification of EVs released in drug tolerant cell lines. (a-c) Cis-tolerant MBGroup3 cells release significantly more EVs in comparison to their matched vehicle cell lines. For detailed description of the cell lines and their identifiers used in this work please refer to methodology section 2.1. Cis resistant & cis treated cell lines are denoted as DT-D458-Cis^DT^, DT-D283-Cis^DT^, DT-HD-MB03-Cis^DT^ and vehicle cell lines are denoted as DT-D458-DMF, DT-D283-DMF, DT-HD-MB03-DMF cell line **(d)** EVs release from Vin-tolerant DAOY cells.

### 3.2. Overcoming EVs mediated cisplatin and vincristine resistance in medulloblastoma

After establishing EV mediated drug resistance in medulloblastoma cell lines, we sought to provide evidence to this effect and investigate whether bioelectronics systems can be used to deliver electrical input to modulate efflux of drug through this pathway. To study the response to external electrical input on the manipulation of intracellular trafficking modulated by sub- cellular entities such as EVs, we used AC (Fig. 3). AC with frequency range of 1 KHz – 5 MHz at a constant potential of 1 V/cm were utilised to study the response. It is worth emphasising that the AC used in this work are not tumour treating fields, as the electrodes were not shielded by dielectric material, and the applied AC does not cause any significant change in the temperature of cell culture medium.^24^ To electrolysis of cell culture medium potentials above 1 V/cm were avoided. For all 4 cell lines treated, metabolic activity only decreased in the drug tolerant line when an AC was applied in the presence of drugs (Fig. 3 a-d and Fig. S1). This effect increased at higher frequencies (Fig. 3 a-d) and was confirmed to be cell death caused enhanced concentration of drug (cis and vin) within the cytoplasm (Fig. 3 e-h) as the drug tolerant cell lines which weren’t treated with Cis/Vin didn’t showed any cell death (Fig. S2).This was in-fitting with previous findings that outer membranes are capacitively coupled to AC at high frequency (KHz-MHz). This leads to membrane electro-permeabilization, which further allows the effect of HF-AC won sub-cellular structures.^25^ Furthermore, no change in the viability of control cells (not drug treated) after the treatment with HF-AC suggests the interaction of HF-AC with membrane bound EVs carrying anticancer drugs.. This could be supported by previous reports in literature where low voltage electric fields lead to disruption of EVs and release of their content. ^26^ Therefore, based on the obtained data it can be concluded that HF-AC could manipulate intracellular drug trafficking by affecting membrane bound EVs, which leads to increased vulnerability of resistant cells towards cis and vin.

**Figure 3.**
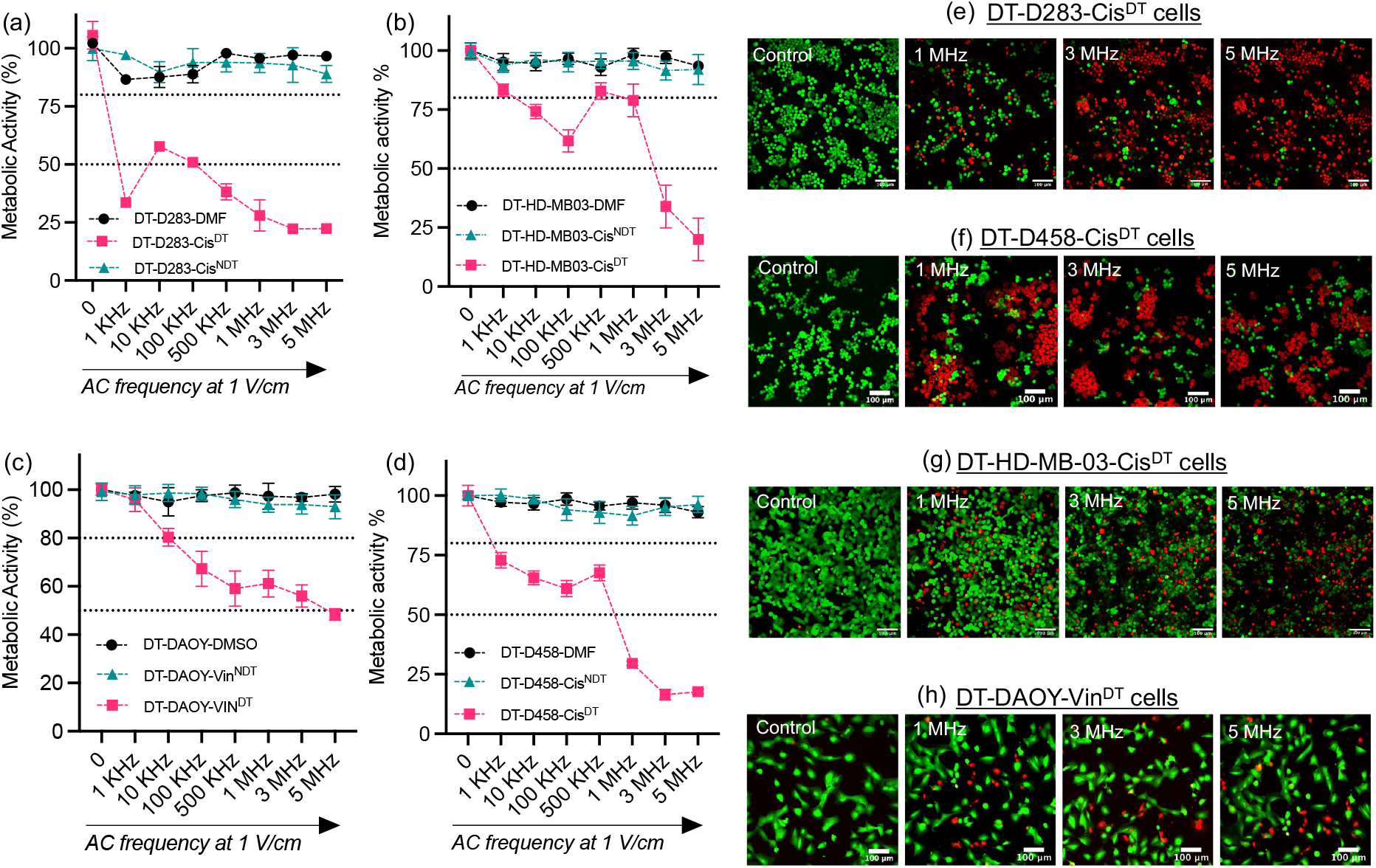
AC-EFs overcomes cisplatin and vincristine resistance in medulloblastoma cells *in vitro*. The cells were stimulated with sinewave AC (1 MHz, 3 MHz, and 5 MHz) using a frequency generator at a potential of 1 V/cm for 30 min followed by a PrestoBlue assay 24 h post electrical stimulation. For detailed description of the cell lines and their identifiers used in this work please refer to methodology section 2.1. **(a-c)** The metabolic activity of cis resistant cells treated with ci (DT-D458-Cis^DT^, DT-D283-Cis^DT^, DT-HD-MB03-Cis^DT^), cis-resistant but non-cis treated cell lines (DT-D458-Cis^NDT^, DT-D283-Cis^NDT^, DT-HD-MB03-Cis^NDT^), and vehicle cell lines (DT-D458-DMF, DT-D283-DMF, DT-HD-MB03-DMF) cell line**. (d)** Vin- resistant cells treated with vin (DT-DAOY-Vin^DT^), vin resistant but non-vin treated cell lines (DT-DAOY-Vin^NDT^), and vehicle (DT-DAOY-DMSO) cell lines. ^NDT^ = no drug treatment, ^DT^ = drug treated. The error bars represent S.E.M of three independent experiment in triplicates. **(e-h)** Live/dead staining of cis and vin resistant medulloblastoma cell lines. The cells were stained with calcein AM (green, live cells) and propidium iodide (red, dead cells) 24 h after stimulation with AC-EFs and imaged using GFP and Texas red filter in a Leica TCS SPE Confocal Microscope. Scale bars = 100 µm.

### 3.3. Endo/lysosomal escape of gold nanoparticles

We next tested our hypothesis that HF-AC affects subcellular structures involved in intracellular trafficking by studying the endosomal escape of GNPs in primary heterogenous brain tumour cell cultures after the treatment with HF-AC (Fig. 4). HF-AC (1-5 MHz at 1 V/cm) was able to induce endosomal escape of GNPs from primary glioma cells isolated from the invasive edge viz. GIN cells (Fig 4. a) and the tumour core viz. GCE cells (Fig. 4 b). No significant toxicity was observed after GNP, AC and GNP + AC treatment (Fig. S3). This was further validated by obtaining Pearson’s correlation coefficient (PCC) which predicted the degree of overlap between green channel (late endosomes) and GNPs (red).^27^ The low values of the PCC obtained from confocal microscopy images shown in Fig 4 a and b, confirmed that application of HF-AC leads to endosomal escape of GNPs. This could be explained based on previous studies that suggest that HF-AC can induce transmembrane potentials which cause transient disruption in cell membrane structures, without causing appreciable toxicity. Importantly, it has been reported that low MHz frequencies can penetrate deep into the cytoplasm to manipulate sub-cellular structures.^28^ Based on these studies, the possibility of GNP escape from endosomes cannot be ruled out however leaking of 100 nm GNPs due to the electro-permeabilization of endosomal membrane requires further studies. Another possible mechanism that could be considered is the behavior of GNPs as electric field transducers.^29^ In this case, the polarization of GNPs in presence of AC could allow them to interact with plasma membrane in a way that facilitates their transport across. However, it is unclear how HF-AC could manipulate endosomal membrane, thus highlighting the need for new investigations on understanding the underlying mechanism such as AC mediated proton sponge effect. In literature there are various reports about the use of external stimuli, such as light and ultrasound, to enhance the cytoplasmic concentration of drugs from polymeric or metallic nanoparticles traversing outside the endosomal compartment.^30,31^ However, it must be emphasized that this effect relies on the properties of the conjugating ligand or peptides that facilitate endosomal escape, but not on the external stimuli. Although conjugation of ligands that enhances endosomal escape has shown great potential, they have been criticized for causing off-set toxicity and reducing the surface coverage for the attachment of targeting moieties.^31, 32^ On the other hand, this work highlights the potential of using external electrical stimuli such as AC, to enhance cytoplasmic concentration of not only small molecular weight drugs but also metallic nanoparticles.

**Figure 4.**
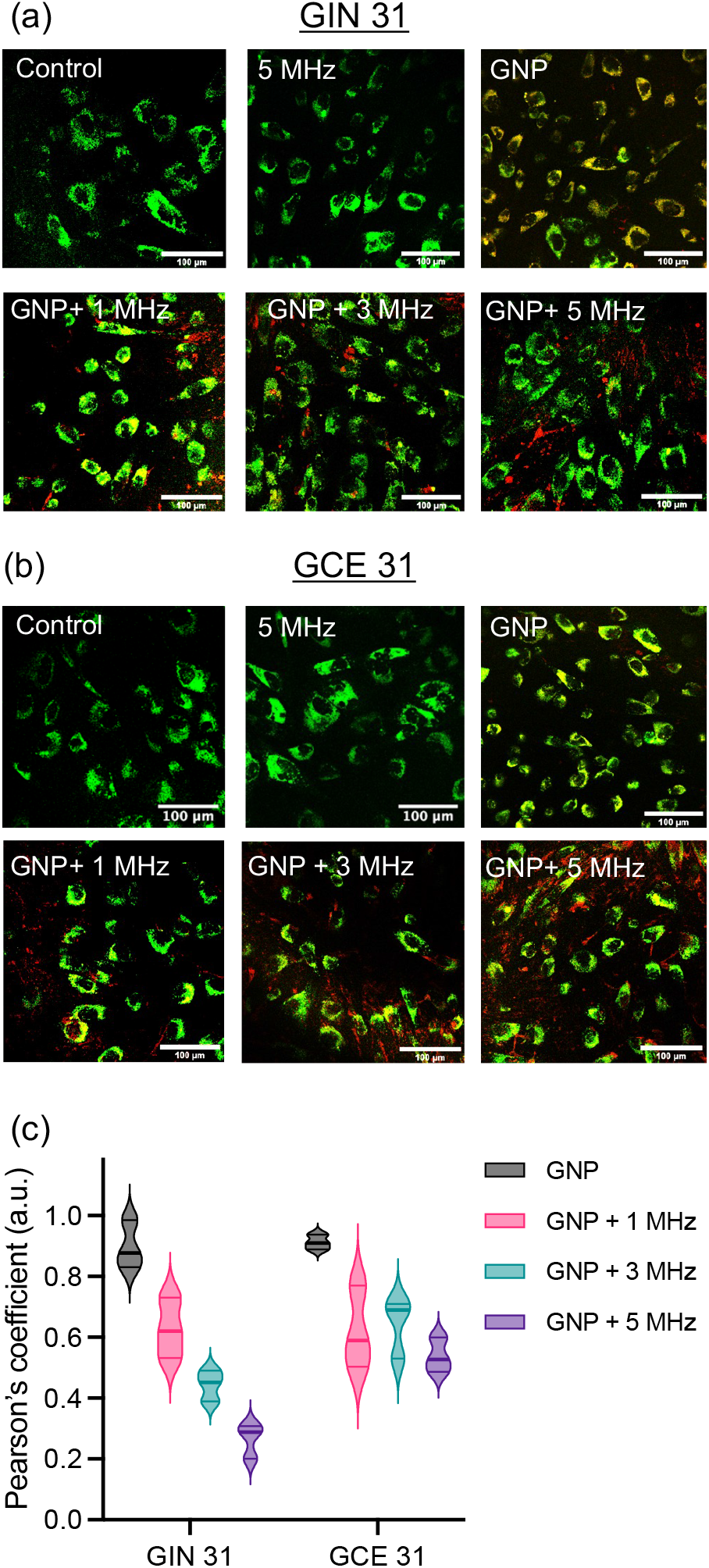
AC mediated endosomal escape of AuNPs in patient derived glioblastoma cells (GIN 31 and GCE 31). Confocal microscopy images to demonstrate endosomal escape of AuNPs (8-hour incubation with cells) in **(a)** GIN 31 and **(b)** GCE 31 cells immediately after treatment with sine wave HF-AC (3 MHz and 5 MHz) at a potential of 1V/cm for 30 min. Cells were stained with late endosome dye (green) and imaged using a Leica confocal microscope with GFP (late endosomes) and Texas-red (AuNPs) filter settings. Scale bar = 100 µm. For detailed description of the cell lines and their identifiers used in this work please refer to methodology section 2.1. To confirm the co-localization of AuNPs at least 50 cells were analyzed. **(c)** Violin plot depicting Pearson’s correlation coefficient (obtained using colocalization plugin in ImageJ) to quantify co-localization of AuNPs (red) with late endosome (green) upon application of EFs. A value of 1 represents perfect co-localization of Texas red (AuNPs) with green channel (late-endosome).

## 4. Conclusions

Our findings support the hypothesis that chemotherapeutic resistance in aggressive brain tumors may be mediated via intracellular trafficking of increased numbers of EVs. Importantly, we have shown that AC can disrupt this EVs mediated trafficking of anticancer drugs out of the cell to enhance vulnerability in drug treated medulloblastoma cells. Furthermore, we showed that HF-AC could enhance the endosomal escape of GNPs in patient-derived GBM cells. Overall, together with ease of HF-AC delivery with no appreciable toxic effects on cells by itself, potentiates the future application of AC in drug delivery to achieve enhanced therapeutic efficacy for better treatment outcomes of cancer.

## 5. Acknowledgements

This work was supported by the Engineering and Physical Sciences Research Council Grant number [EP/R004072/1].

## 6. Authors Contribution

**Akhil Jain:** Conceptualization, Methodology, Validation, Investigation, Writing - original draft, Visualization, Writing - review & editing, Formal analysis, Supervision, Funding acquisition. **Philippa Wade:** Investigation, Writing - original draft, Visualization, Writing - review & editing, Formal analysis. **Snow Stolnik:** Supervision, Writing - review & editing. **Alistair N Hume**: Supervision, Writing - review & editing. **Ian D. Kerr:** Supervision, Writing - review & editing. **Beth Coyle:** Conceptualization, Methodology, Validation, Writing - original draft, Writing - review & editing, Supervision, Funding acquisition. **Frankie Rawson:** Conceptualization, Methodology, Validation, Writing - original draft, Writing - review & editing, Supervision, Funding acquisition.

## 8. Supporting Information

The data related to live dead staining of medulloblastoma cells at frequencies below <1 MHz (Fig. S1), live dead images of vehicle cell lines (Fig. S2), and biocompatibility of GNPs in primary GBM cells (Fig. S3).

## 9. Data Availability

All the data will be available to the readers at free of cost at https://nottingham.rdmc.ac.uk.

---

**Figure S1.**
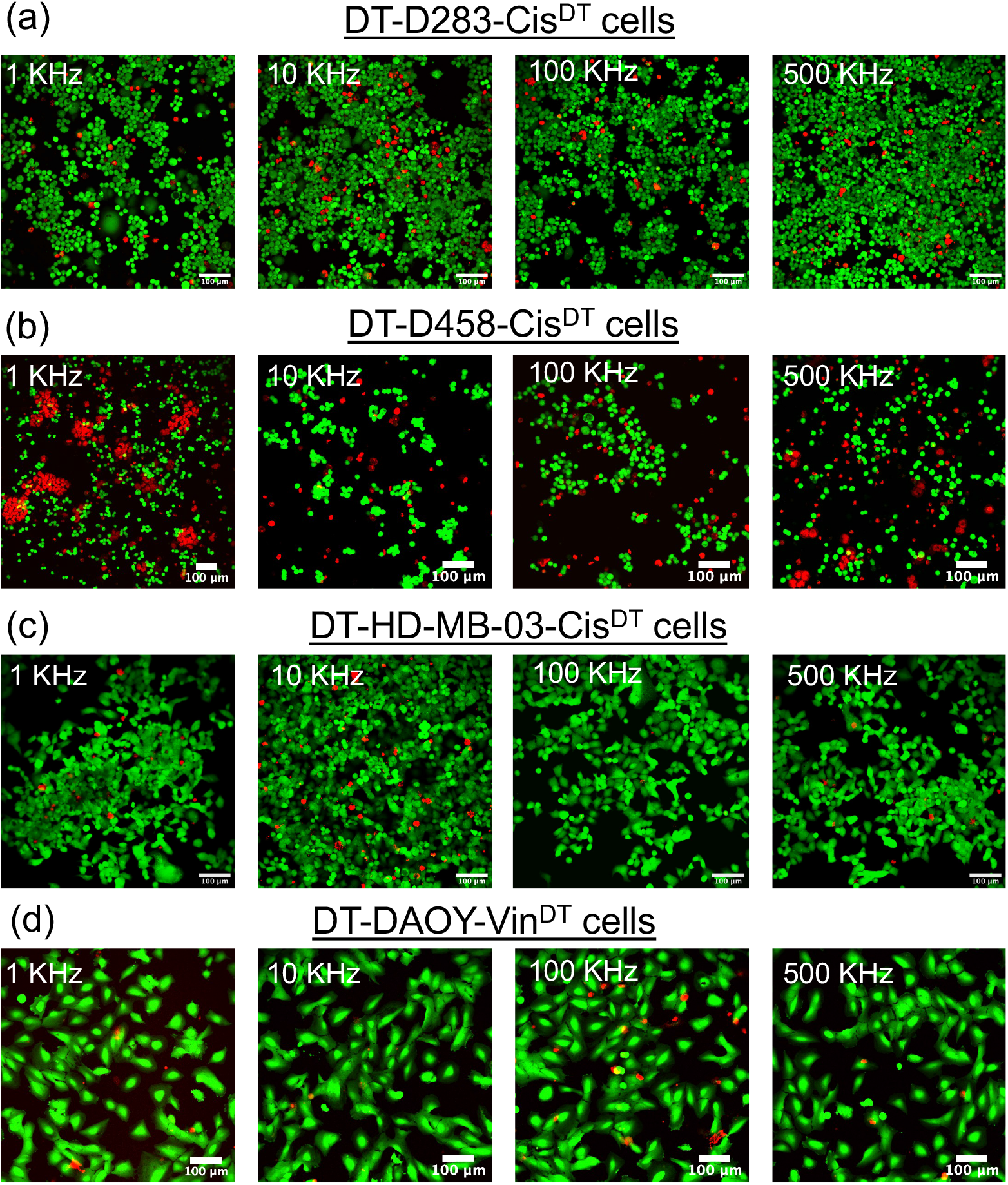
– AC-EFs overcomes cis and vin resistance in medulloblastoma cells *in vitro*. The cells were stimulated with square wave AC-EFs (1 KHz, 10 KHz, 100 KHz, and 500 KHz) using a frequency generator at a potential of 1V/cm for 30 min. Live/dead staining of cis and vin resistant medulloblastoma cell lines. The cells were stained with calcein AM (green, live cells) and propidium iodide (red, dead cells) 24 h after stimulation with AC-EFs and imaged using GFP and Texas red filter in a Leica TCS SPE Confocal Microscope. Scale bars = 100 µm.

**Figure S2.**
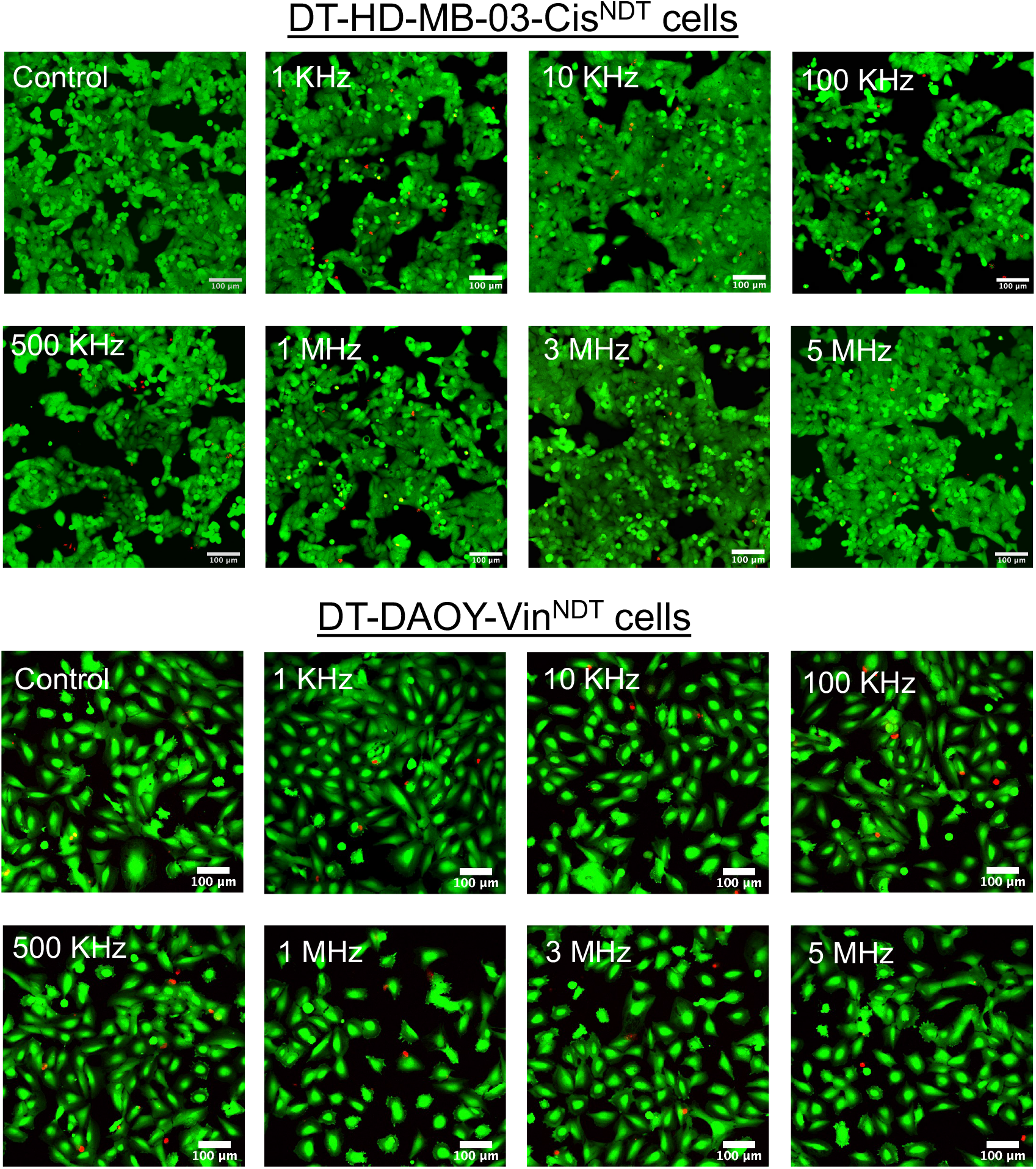
Live/dead staining of vehicle medulloblastoma cell lines. The cells were stained with calcein AM (green, live cells) and propidium iodide (red, dead cells) 24 h after stimulation with AC-EFs and imaged using GFP and Texas red filter in a Leica TCS SPE Confocal Microscope. Scale bars = 100 µm.

**Figure S3.**
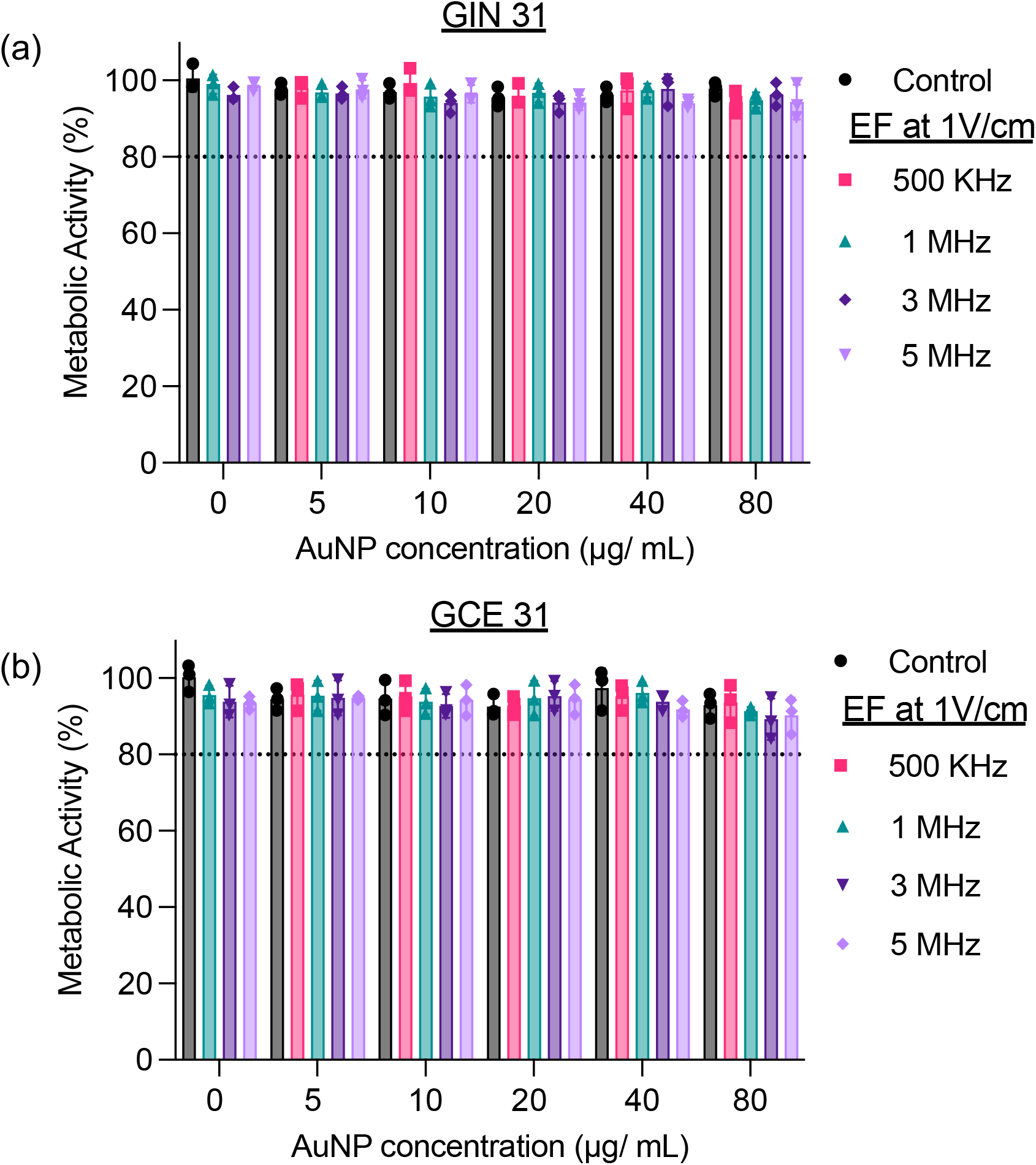
Biocompatibility of AuNPs on GIN 31 and GCE 31 in presence of AC-EFs. The cells were treated with AuNPs for 8 h before stimulation with square wave AC-EFs (1 KHz, 10 KHz, 100 KHz, and 500 KHz) using a frequency generator at a potential of 1V/cm for 30 min. The metabolic activity of cells was determined 24 hours after stimulation with AC-EFs using PrestoBlue assay. The error bars represent the S.E.M. from a triplicate experiment repeated thrice.

